# QM/MM simulations of EFGR with afatinib reveal the role of the *β*-dimethylaminomethyl substitution

**DOI:** 10.1101/2024.02.18.580887

**Authors:** Shuhua Ma, Heeral Patel, Craig A. Peeples, Jana Shen

## Abstract

Acrylamides are the most commonly used warheads of targeted covalent inhibitors (TCIs) directed at cysteines; however, the reaction mechanisms of acrylamides in proteins remain controversial, particularly for those involving protonated or unreactive cysteines. Using the combined semiempirical quantum mechanics (QM)/molecular mechanics (MM) free energy simulations, we investigated the reaction between afatinib, the first TCI drug for cancer treatment, and Cys797 in the EGFR kinase. Afatinib contains a *β*-dimethylaminomethyl (*β*-DMAM) substitution which has been shown to enhance the intrinsic reactivity and potency against EGFR for related inhibitors. Two hypothesized reaction mechanisms were tested. Our data suggest that Cys797 becomes deprotonated in the presence of afatinib and the reaction proceeds via a classical Michael addition mechanism, with Asp800 stabilizing the ion-pair reactant state *β*-DMAM^+^/C797^−^ and the transition state of the nucleophilic attack. Our work elucidates an important structure-activity relationship of acrylamides in proteins.

## INTRODUCTION

Kinases are enzymes that catalyze the phosphoryl transfer reactions; dysregulation of kinases has been linked to a variety of human diseases especially cancer.^1^ In 2013 the first targeted covalent inhibitor (TCI), afatinib, which inhibits a receptor kinase called epidermal growth factor receptor (EGFR, Figure 1a and 1b), obtained the FDA approval for treatment of non-small-cell lung (NSCL) cancer.^2^ Over the last decade, FDA approvals of TCIs of various kinases have been steadily increasing.^3^ A TCI is comprised of a reversible binding motif and an electrophilic functional group (also called the warhead) that forms a chemical bond with a nucleophilic amino acid (typically cysteine) in the target protein. Understanding of the chemical reaction mechanism of TCIs in the protein environment is important for the rational design of novel inhibitors and improvement of existing ones.

**Figure 1.**
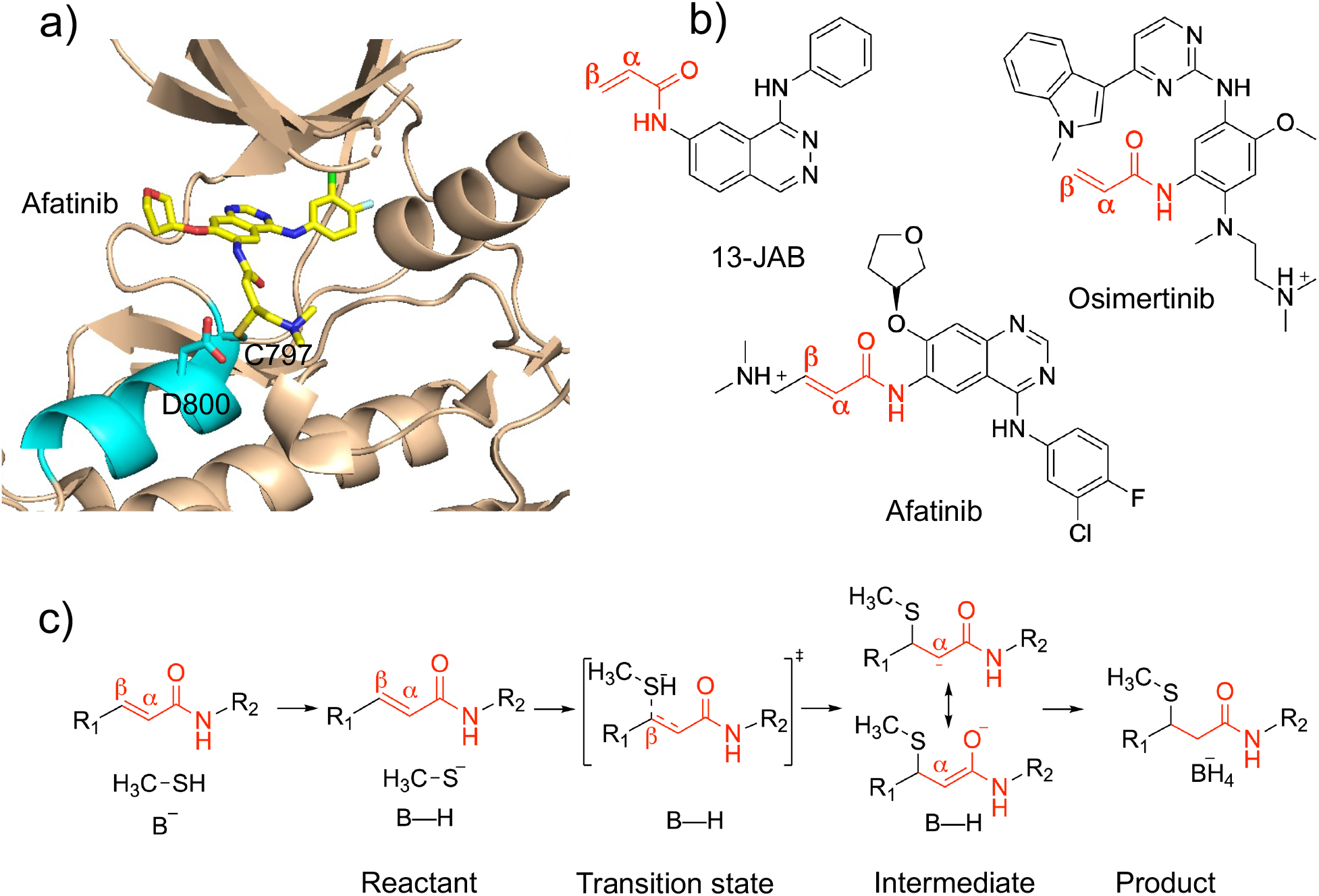
The TCIs of EGFR studied by QM/MM simulations and the thiol-Michael addition reaction mechanism in solution. **a)** The X-ray cocrystal structure of EGFR in covalent complex with afatinib (PDB ID: 4G5J). ^16^ The *α*D helix is highlighted in cyan. The covalently modified Cys797 and the nearby Asp800 which can be a potential general base are labeled. **b)** Chemical structures of afatinib and two previously studied ^17,18^ TCIs of EGFR. **c)** The base-assisted mechanism of Michael addition between acrylamide and thiol (shown here as a methanethiol) in solution. B-H represents a general base. The resonance of the carbanion/enolate intermediate is shown.

So far, all FDA-approved TCIs contain an acrylamide warhead, which is a type of activated olefin capable of forming a covalent bond with cysteine’s thiol group through Michael addition. Based on the solution-phase experiments^4–6^ and density functional theory (DFT) calculations,^7–11^ it is believed that the rate-limiting step of a thiol-Michael addition is the nucleophilic attack of a deprotonated thiolate on the electron deficient *β*-carbon of the olefin, resulting in a carbanion intermediate which reprotonates to yield the adduct (Figure 1c). In the absence of a deprotonated thiolate, a Michael reaction can be initiated, in which a nearby base abstracts the proton from the thiol group, and the same proton is later transferred to protonate the carbanion intermediate^12^ (Figure 1c). Our recent studies of various kinases^13–15^ based on the continuous constant pH molecular dynamics (CpHMD) titration simulations found that many cysteines targeted by the TCIs approved by the FDA or in clinical stages are indeed reactive, as defined by having a deprotonated population greater than 10% at physiological pH 7.4.

Of particular interest is the so-called front-pocket cysteine located at the N-terminal end of the *α*D helix in 11 kinases including EGFR. The front pocket cysteines of all these kinases have been targeted by the TCIs, making it the most popular site of covalent modification for kinases.^14^ The CpHMD simulations suggested that the front-pocket cysteine is reactive in the apo state of 7 kinases, e.g., Cys481 in BTK; however, it is unreactive in EGFR (Cys797) and the related ERBB2 and ERBB4 as well as BLK, due to a negatively charged Asp or Glu sidechain at the Cys+3 position (Asp800 in EGFR).^14^ Considering the proximity of the carboxylate to the cysteine thiol group in the cocrystal structures, the Cys+3 Asp or Glu may serve as a general base to abstract the proton from the front-pocket cysteine and thereby initiating the Michael reaction.^14^

The reaction of two TCIs with the front-pocket Cys797 in EGFR has been studied by the combined quantum mechanics/molecular mechanics (QM/MM) simulations which started from the protonated Cys797. Capoferri et al.^17^ investigated the reaction mechanism of 13-JAB (Figure 1b) with EGFR using the semiempirical DFTB/MM potential. Using the same method, Callegari et al.^18^ examined the effect of EGFR mutations on the reaction barrier with osimertinib (Figure 1b). Surprisingly, both studies^17,18^ suggested that the nucleophilic attack of the deprotonated Cys797 is concerted with the proton transfer from the protonated Asp800 without a stable intermediate. This direct addition mechanism contradicts the stepwise Michael addition mechanism suggested by solution experiments^4–6,19^ and QM calculations of thiol additions of activated olefins;^8,9,11,12,20,21^ however, the contradiction might be due to the difference between solution and the protein environment.

More recently, two QM/MM studies^22,23^ examined the reaction mechanism of the acrylamide TCIs with the front-pocket cysteine Cys481 in BTK, which was found to be deprotonated in the CpHMD simulations.^14^ The QM/MM simulations employing the density functional theory (DFT) QM description of a truncated *α*-cyano acrylamide inhibitor and starting from a deprotonated Cys797 confirmed the Michael reaction mechanism involving an intermediate.^23^ In contrast, the QM/MM simulations using the DFTB QM description of BTK and protonated Cys481 suggested an alternative mechanism which involves first a proton transfer from Cys481 to the carbonyl group and the subsequent nucleophilic attack is not rate limiting.^22^ The different conclusions from the above four studies^17,18,22,23^ suggest that acrylamide-cysteine reactions in the protein environment remain poorly understood.

In the present work, we use afatinib’s reaction with the front-pocket Cys797 in EGFR as a model system to understand the mechanism of the covalent addition of acrylamides to an unreactive cysteine. Afatinib is an ATP-competitive anilinoquinazoline derivative; its acrylamide warhead has a dimethylaminomethyl (DMAM) substitution on the *β*-carbon (Figure 1b). Two early experimental studies found that *β*-DMAM substitution enhances the reactivity (i.e., rate constant) of quinazoline-based acrylamides (afatinib analogs) against glutathione in solution and the inhibitory activity against EGFR.^24,25^ The latter was demonstrated by more than 20 times decrease of the IC50 value in the en-zyme assay and 6–550 times decrease in three cell-based assays.^24^ We note, a more recent study^26^ using two analogs of afatinib showed a reversed but much smaller effect of *β*-DMAM substitution in the EGFR mutant L858R/T790M. Specifically, the rate constant (*k*_inact_) is decreased by half and the IC50 value in one cell assay is increased by 7 times with the *β*-substitution.^26^

A most recent experiment^10^ demonstrated that the glutathione half-life of the *β*-methylamino substituted acrylamides decreases with the increasing p*K* _a_ value (from 6 to 9) of the amino substituent, implying that a positively charged *β* substitution increases the Michael reaction rate. To explain this, our recent QM calculations of model reactions in solution (implicit solvent model)^11^ found that a charged *β*-DMAM group lowers the reaction barrier of nucleophilic attack through an induction effect by increasing the accumulated charge on the *α*-carbon in the carbanion intermediate state. However, in the protein environment, the reaction mechanism of *β*-DMAM substituted inhibitors may be different from that in solution, especially in the presence of a protonated cysteine. Indeed, while studying the structure-activity relationship of acetylenic thienopyrimidine compounds, Wood et al.^27^ hypothesized that the amino substituent at the C*β* position is the general base that deprotonates Cys797 before nucleophilic attack.

Inspired by the aforementioned work and hypotheses, here we conducted the semiempirical QM/MM free energy simulations to investigate the afatinib reaction with Cys797 in EGFR. We aim to address the following questions that have generated controversies in the previous QM/MM studies of EGFR and BTK reactions with acrylamide inhibitors.^17,18,22,23^ 1) How does Cys797 become deprotonated? 2) Is the nucleophilic attack rate limiting? 3) Is there an enolate/carbanion intermediate? Considering that Cys797 is protonated in the absence of the inhibitor,^14^ we investigated two possible mechanisms, which differ in how Cys797 becomes deprotonated, a requirement for nucleophilic attack (Scheme 1). Specifically, Mechanism 1 tested the hypoth-esis that a neural *β*-DMAM group serves as a general base/acid^27^ to abstract the proton from Cys797, while Mechanism 2 tested the hypothesis that the nearby Asp800 serves as a general base/acid.^15^ The free energy simulations of Mechanism 1 found that the ion-pair reactant state *β*-DMAM(+)/C797(-) is significantly more stable than the neutral counterpart, and the subsequent nucleophilic attack has a barrier similar to the experimental estimate. However, if Cys797 deprotonates by donating the proton to the nearby Asp800 (Mechanism 2), the resulting nucleophilic attack has a significantly higher barrier. We discuss the reaction paths and the nature of the intermediate states as well as their stabilization.

## RESULTS and DISCUSSION

### Two possible reaction mechanisms

We investigated two possible reaction mechanisms of afatinib with Cys797 in EGFR (Scheme 1). Mechanism 1 begins with the first proton-transfer step (PT-1), in which Cys797 thiol transfers a proton to the *β*-DMAM nitrogen of the acrylamide, initiating the nucleophilic attack (NA) of the thiolate on the *β*-carbon followed by the second proton-transfer step (PT-2), whereby the proton is transferred from the *β*-DMAM nitrogen to the *α*-carbon, resulting in the thioether adduct product (Scheme 1, top). In contrast, Mechanism 2 begins with a proton transfer from the cysteine thiol to Asp800 in EGFR (PT-1), following the NA step, the proton on Asp800 is transferred to the *α*-carbon (PT-2, Scheme 1, bottom). PT-1 and NA, and similarly also NA and PT-2, may be stepwise or concerted, which can be distinguished through QM/MM simulations.

### Summary of the PDDG-PM3/MM simulations

The simulations were initiated from the cocrystal structure of EGFR in complex with the covalently bound afatinib (PDB ID: 4G5J).^16^ For Mechanism 1, the QM region in-cludes the sidechain of Cys797 and the entire inhibitor (afatinib), while the C*α* atom of Cys797 is the QM/MM boundary atom (Figure 2a and b). For comparison, the simulations of the reaction steps in solution were also performed, the QM region in solution includes methanethiol and afatinib. For Mechanism 2, the QM region includes the the entire inhibitor (afatinib) and the sidechains of Cys797 and Asp800, and the C*α* atoms of Cys797 and Asp800 are the QM/MM boundary atoms. Due to the large number of atoms in the QM region (65 atoms for Mechanism 1 and 73 atoms for Mechanism 2), the semiempirical Pairwise Distance Directed Gaussian modified PM3 method (PDDG-PM3)^28,29^ was used (details see Methods).

**Figure 2.**
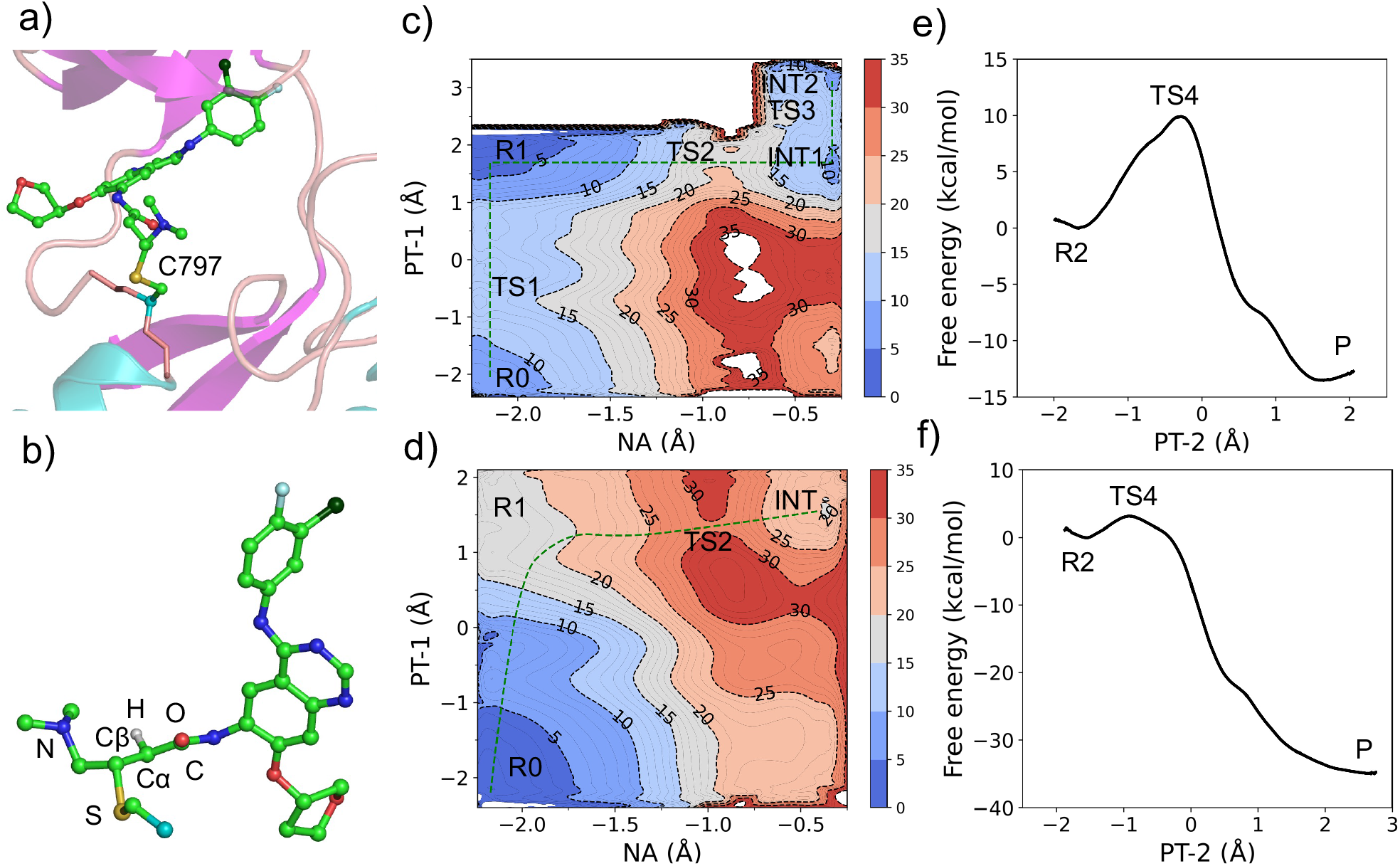
QM/MM free energy simulations of the covalent bond formation between afatinib and Cys797 in EGFR and the model reaction in solution. **a)** Partition of the QM (stick model) and MM (cartoon model) regions in the combined QM/MM simulations. Water (hidden) is in the MM region. **b)** A zoomed-in view of the QM region. Atoms involved in the chemical steps are labeled. Except for the reactive one, hydrogens are hidden for clarity. **c, d)** Calculated free energy surface along the reaction coordinates of the NA 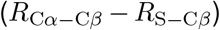 and PT-1 (*R*_S−H_ *− R*_H−N_) steps in EGFR (c) and model reaction in solution (d) according to Mechanism 1. *R* is the distance of the breaking or forming bond. **e, f)** Calculated free energy change along the reaction coordinate of PT-2 (*R*_N−H_ − *R*_H−Cα_) in EGFR (e) and model reaction in solution (f) according to Mechanism 1. The state labels are explained in the main text and correspond to those in Scheme 2.

**Scheme 1:**
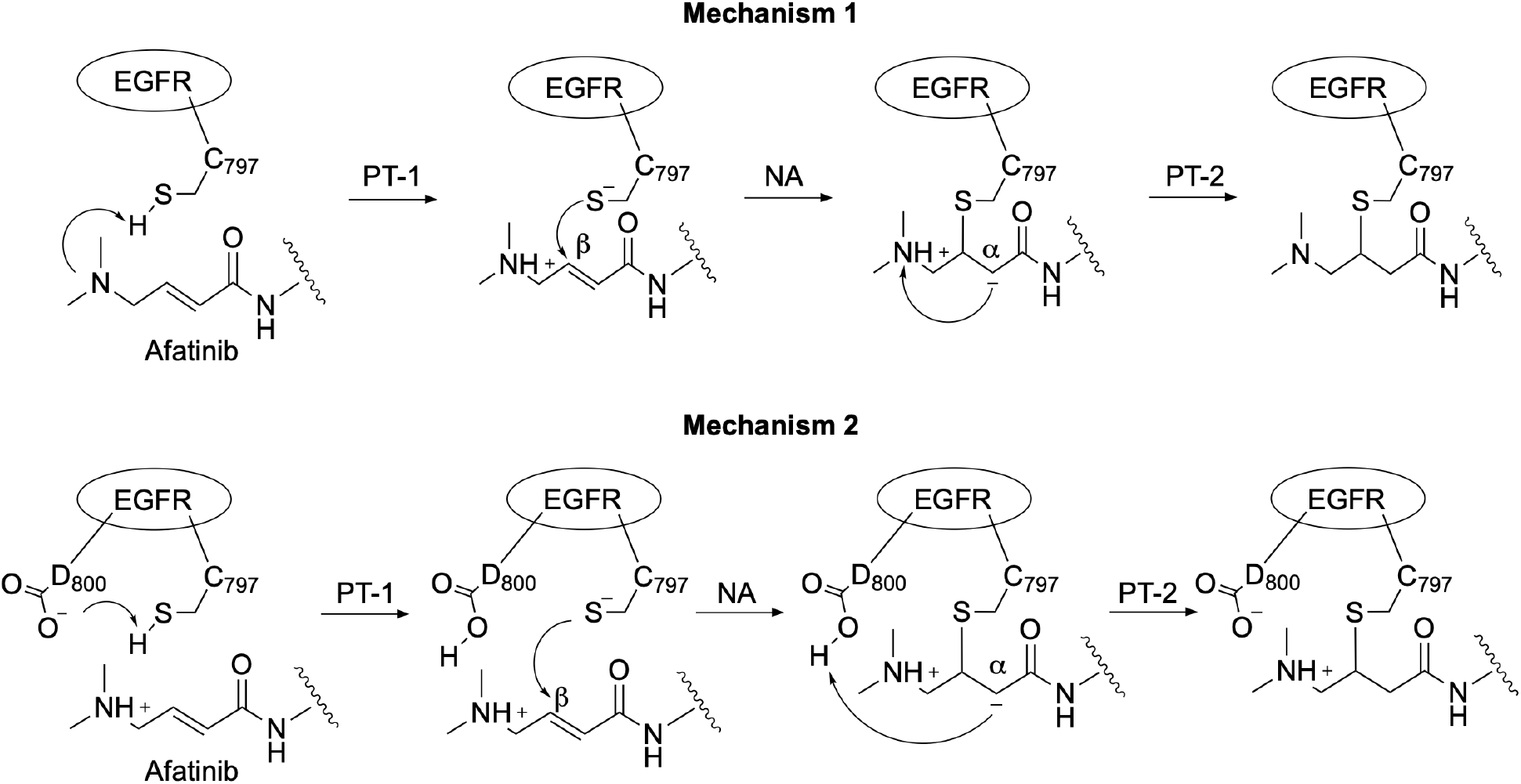
Two possible reaction mechanisms of afatinib covalent addition with Cys797 of EGFR investigated in this work. The Mechanism 1 and 2 differ in that either the *β*-DMAM group of the acry-lamide warhead or the nearby Asp800 acts as a general base to abstract the proton from the otherwise unreactive Cys797 (PT-1) before nucleophilic attack (NA) of thiolate on C*β* followed by the reprotonation of C*α* atom (PT-2). Note, other possible variants of Mechanism 1 are not shown, e.g., 1) the product of PT-2 in Mechanism 1 becomes protonated by solution water; 2) instead of PT-2, the carbanion can be protonated by solution water.

### PT-1/NA steps of Mechanism 1 are step-wise in EGFR but concerted in solution

To investigate the multi-step reaction Mechanism 1, we first performed two-dimensional QM/MM free energy simulations along the re-action coordinates that represent the proton transfer from the cysteine thiol to the nitrogen of the *β*-DMAM group on afatinib (PT-1) and the nucleophilic attack (NA) from the cysteine sulfur to the C*β* of afatinib (Figure 2c and d). For PT-1, the difference between the thiol sulfur-to-hydrogen and *β*-DMAM hydrogen-to-nitrogen distances was used as the reaction coordinate. For NA, the difference between the C*α*-to-C*β* and sulfur-to-C*β* distances was used as the reaction coordinate. The resulting free energy surface (Figure 2c) shows that the ion-pair state (R1) with a deprotonated thiolate and charged *β*-DMAM group is more stable than the neutral state (R0), which means that Cys797 exists as thiolate and *β*-DMAM group is charged in EGFR upon afatinib binding. Thus, the covalent bond formation process starts from the R1 state, from which the nucleophilic attack occurs and has a free energy barrier of about 20 kcal/mol. The nucleophilic attack results in an intermediate state (INT1), which then underwent a conforma-tional change to form a slightly more stable INT2, the two intermediate states are separated by a small free energy barrier, and INT2 state serves as the reactant state (R2) for the PT-2 step.

As a comparison and reference, we also conducted the QM/MM free energy simulations of the same PT-1 and NA reactions of Mechanism 1 in solution. From the free energy surface, it is clear that methanethiol and afatinib exist in the neutral state, as the ion-pair state (R1) has a much higher free energy (close to 16 kcal/mol than the neutral state (R0, Figure 2d). This result is consistent with the high p*K* _a_ of 10.33 for methanethiol in water.^30^ The free energy surface shows that the nucleophilic attack and proton transfer from methanethiol and *β*-DMAM is concerted with a much higher free energy barrier than that in EGFR (Figure 2d). The resultant carbanion/enolate intermediate (INT1) has a much higher free energy relative to the reactant state R0.

### PT-2 step of Mechanism 1 in EGFR has a higher barrier but is less exergonic compared to that in solution

We next performed the one-dimensional free energy simulations to investigate proton transfer (PT-2) from the *β*-DMAM nitrogen to the C*α* atom of afatinib in both EGFR and solution (Figure 2e and f). For PT-2, the difference between the *β*-DMAM nitrogen-to-hydrogen and hydrogen-to-C*α* distances was used as a reaction coordinate. The free energy of activation in EGFR is significantly higher than in the model reaction solution. However, the reaction is significantly less exergonic in EGFR as compared to solution.

### Analysis of the reaction profiles of Mechanism 1

The free energy profile of Mechanism 1 in EGFR (red) and solution (black) is summarized in Scheme 2. Note, a DFT-based correction is applied (Supplemental methods). In EGFR, the covalent bond formation between afatinib and Cys797 involves three steps, PT-1, NA, and PT-2. The ac-tivation barriers for PT-1 (from R0 and R1), NA (from R1 to INT1), and PT-2 (from R2 to P) steps are 7.3 kcal/mol, 19.5 kcal/mol, and 9.9 kcal/mol, respectively. From the INT1 intermediate a conformational rearrangement of the *β*-DMAM group occurs with a free energy barrier of 3.9 kcal/mol, leading to the slightly more stable INT2 intermediate, which is 8.3 kcal/mol higher in free energy than the reactant R1 state. By contrast, the afa-tinib reaction with methylthiol in solution proceeds from the R0 state through concerted PT-1 and NA steps, however the NA step is rate-limiting with a free energy barrier of 29.8 kcal/mol, which is about 10 kcal/mol higher than the NA step in EGFR. However, the PT-2 step in water has a much lower free energy barrier of 3.2 kcal/mol, as compared to 9.9 kcal/mol in EGFR. These data show that in both EGFR and solution, the rate-limiting step is the NA step. Note, the former (19.5 kcal/mol in EGFR) is in good agreement with the free energy of activation of 21.6 kcal/mol converted from the measured rate constant of 9 *×* 10^−4^s^−1^ for the covalent inhibition of EGFR by afatinib.^26^ The reaction free energy in EGFR (from R1 to P) is -3.2 kcal/mol, which is much smaller in magnitude than the corresponding reaction free energy (from R0 to P) of -9.8 kcal/mol in solution. The reaction free energy of the PT-2 step (from R2 to P) in EGFR is -11.5 kcal/mol, which is also much smaller in magnitude than the corresponding reaction free energy of -29.5 kcal/mol in solution.

### Comparison to the less likely Mechanism2

Now we turn to the afatinib reaction in EGFR according to Mechanism 2, whereby the nearby Asp800 acts as a general base to deprotonate Cys797 (PT-1), followed by NA and reprotonation of C*α* on afatinib (PT-2). We note that the reaction here starts from the charged *β*-DMAM group, because in solution *β*-DMAM is charged^11^ and in the co-crystal structure (PDB 4G5J^16^), the *β*-DMAM nitrogen is only 4.5 Å away from the nearest car-boxylate oxygen of Asp800. The individual PT-1, NA, and PT-2 steps have free energy barriers of 11.7, 23.3, and 7.1 kcal/mol, respectively (Figure S8). The carbanion/enolate intermediate state is 15.4 kcal/mol higher in free energy than the reactant state (R2, thiolate^−^/Asp800^H^). Overall, the rate-liming NA step has a free energy barrier of 31.3 kcal/mol, which is much higher than that in Mechanism 1, we conclude that Mechanism 1 is a more likely reaction path.

The free energy profiles of the afatinib reaction in both EGFR and solution demonstrate a classic Michael addition in which the NA step is rate-limiting and a stable intermediate is formed before reprotonation to the thiol adduct state. This is consistent with Rowley and coworker’s QM/MM simulations of ibrutinib reaction with BTK,^23^ although their simulations started from a deprotonated thiolate state.

### Formation of a carbanion/enolate intermediate

To further understand the reaction Mechanism 1, we examined the structure changes near the reaction center along the reaction path using the configurations from the free energy simulations (details see Supplemental Methods). The selected distances and Mulliken atomic partial charges of the reaction center along the reaction path are given in Table S6 and S7. Of particular interest is the nature of the intermediate state INT2. In the literature, it is often stated that the Michael addition of acrylamides give an enolate which formally has the negative charge localized on the oxygen atom.^12,23,31^ Our QM/MM simulations in both EGFR and solution demonstrated that the negative charge of the intermediate resulting from the NA step is delocalized, i.e., it is in a carbanion/enolate resonance. In the INT2 state of EGFR (Figure 3), the S−Cβ bond is formed (distance of 1.85 Å as compared to 3.36 Å in the R1 state), and the proton re-mains bonded to the *β*-DMAM nitrogen (N–H distance of 1.01 Å). The C*β*−Cα distance is elongated to 1.47 Å from the C**−**C bond dis-tance of 1.34 Å in the R1 state, although still slightly shorter than the C−C bond distance of 1.54 Å in the product state.

**Figure 3.**
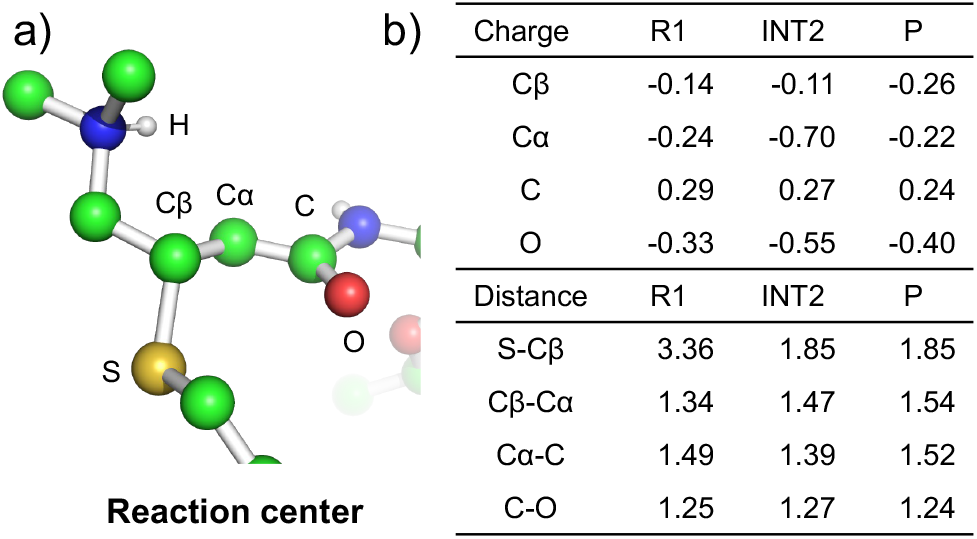
Changes in the bond lengths and atomic partial charges in the reaction center between the reactant, intermediate, and product state. a) The geometry of the reaction center in the INT2 state. Nonpolar hydrogens are hidden. Both afatinib and cysteine are cropped, and the protein (EGFR) as well as water are hidden. Relevant atoms are labeled. b) Relevant bond lengths (Å) and Mulliken partial charges (a.u.) in R1 (before NA), INT2 (carbanion/enolate), and P states.

An ideal enolate has a C*α***−**C bond and a C−O bond, whereas an ideal carbanion has a C*α*−C bond and a C**−**O bond. The INT2 state shows that the C*α*−C and C−O distances are 1.39 and 1.27 Å, respectively, which are shorter than the respective C−C and C−O single bond lengths of 1.54 and 1.43 Å, but longer than the respective C**−**C and C**−**O double bond lengths of 1.33 and 1.23 Å. In so-lution, the aforementioned distances are identical to the values in EGFR. Thus, our QM/MM simulations suggest that the intermediate of the NA step has both the ideal carbanion and enolate characters, which is further demonstrated by the Mulliken charges (Supplemental Table S7). In INT2 of EGFR, the negative charge is distributed to both C*α* (−0.7) and O (−0.55) atoms, with more negative charge on C*α*. In solution, the negative charge is distributed slightly more onto the C*α* atom, with a charge of -0.73 as compared to -0.50 on the O atom.

### Asp800 stabilizes the ion-pair reactant state

To identify critical residues in EGFR that contribute to the formation of the ion-pair state R1 and the final adduct state, we performed the interaction energy decomposition analysis.^32,33^ For each selected residue, we calculated its QM/MM interaction energy with the QM region, which is defined as the difference in the QM energy before and after annihilation of the MM charges on the residue. This calculation was performed for all the trajectory snapshots representing the R0, R1, and P states, and for each state, the interaction energies were averaged for each residue (Supplemental methods for more details). Thus, the interaction energy difference between two states represents the residue-specific contribution to the stabilization or destabilization of the state. Figure 4 shows the residue-based interaction energies for stabilizing (or destabilizing) the R1 state (relative to R0 state) or the P state (relative to R1 state). Three residues make greater than 1 kcal/mol contributions to the stabilization or destabilization of the R1 state relative to the R0 state. Arg803 and Asp855 destabilize the R1 state by 1.2 kcal/mol and 2.5 kcal/mol, respectively, whereas Asp800 stabilizes the R1 state by 8.3 kcal/mol, making it a major contributor to the PT-1 step. Since Asp800 is absent in solution, this may explain why R1 does not occur in solution, as it has much higher free energy (Figure 2d). Examination of the trajectory snapshots revealed that the stabilizing interaction of Asp800 with afa-tinib arises from the electrostatic interaction (a weak hydrogen bond) between the carboxylate oxygen of Asp800 and the *β*-DMAM nitrogen. The distance between the reactive hydrogen of *β*-DMAM and the nearest carboxylate oxygen is *∼*2.9 Å in the R1 state and it reduces to *∼*2.6 Å in the TS2 state. Thus, Asp800 not only stabilizes the ion-pair complex but also lower the free energy barrier of the NA step. The latter effect is missing in solution, leading to a much higher barrier. In the product state, however, Asp800 contributes over 8 kcal/mol destabilization energy with respect to the R1 state, which may explain the less favorable reaction free energy in EGFR as compared to solution (Figure 2).

**Figure 4.**
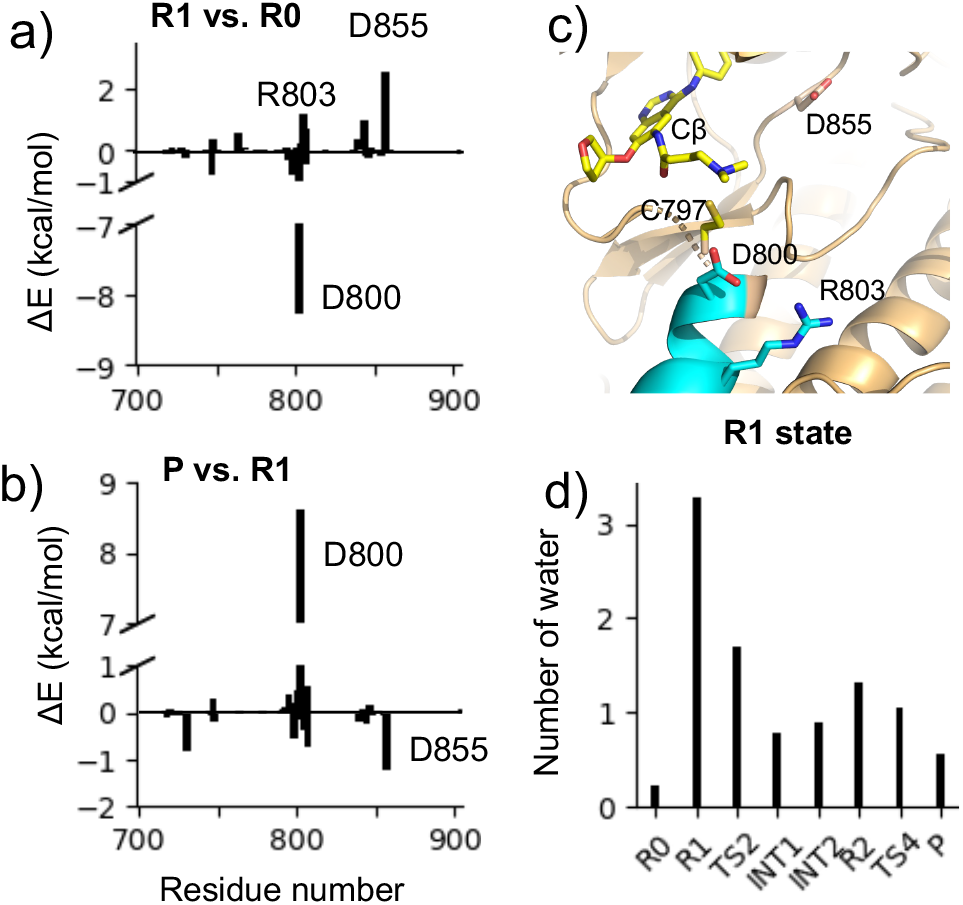
Interaction energy decomposition and hydration analysis. **a**,**b)** Residue-based electrostatic contributions to the QM/MM energy change of the R1 vs. R0 state (a) and the P vs. R1 state (b). Residues with contributions greater than 1.0 kcal/mol are labeled. **c)** A zoomed-in view of the locations of the three residues making the largest electrostatic energy contributions shown in a) and b). **d)** The average number of water within 3.5 Å of the Cys797 sulfur in different states along the reaction path.

**Scheme 2:**
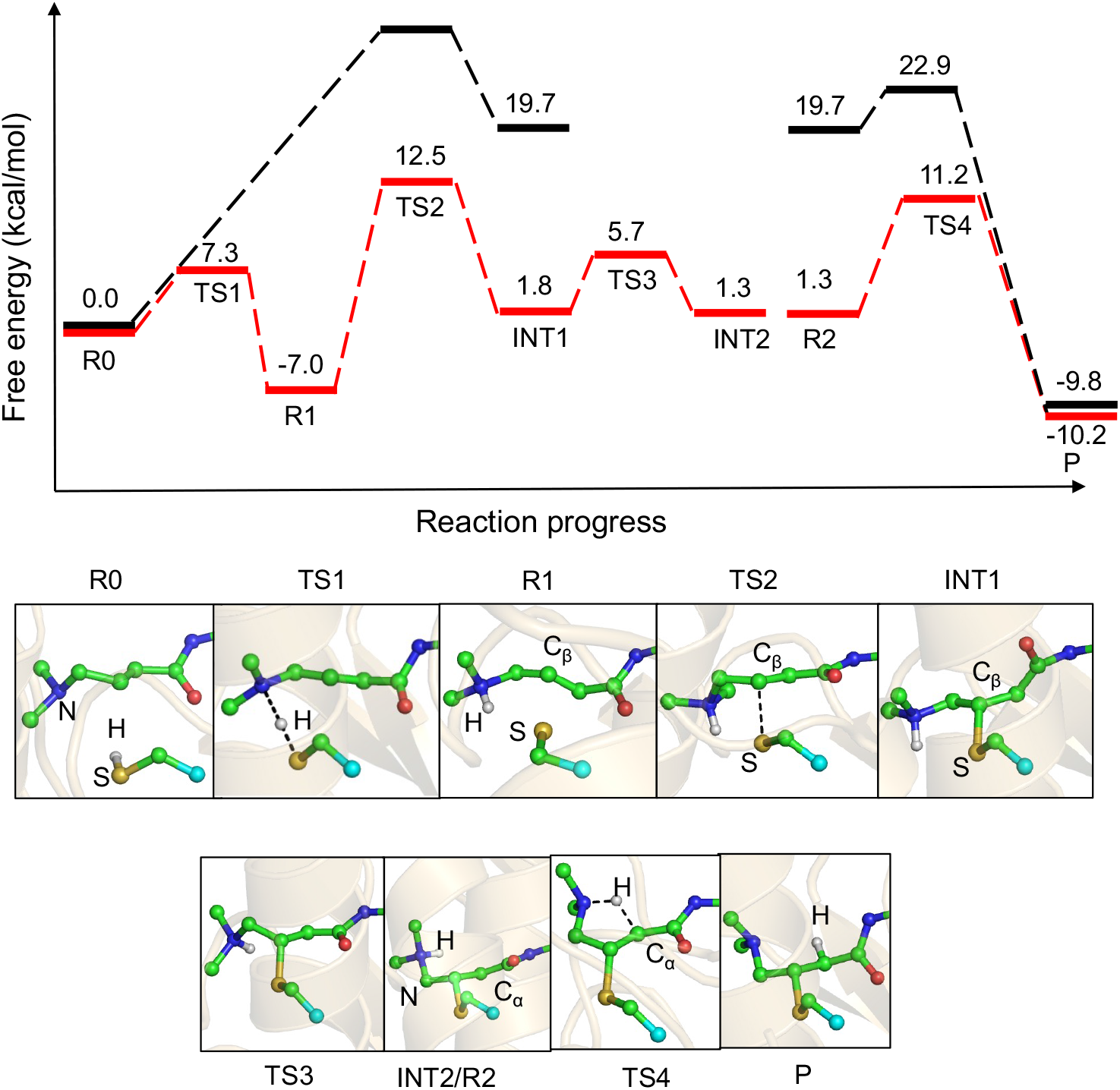
Calculated free energy profile along the reaction path in EGFR (red) and in aqueous solution (black) according to Mechanism 1. The relative free energies with the DFT corrections are given. The representative snapshots of the states along the reaction path in EGFR are shown. The states correspond to those in Figure 2c and d. R0: reactant state with neutral Cys797 (or methanethiol in solution) and *β*-DMAM substituted acrylamide. R1: reactant state (ion-pair) with the charged Cys797 thiolate (or methylthiolate in solution) and *β*-DMAM^+^ substituted acrylamide. TS1: transition state for PT-1 (proton transfer from the thiol to the *β*-DMAM group). TS2: transition state for the NA step. INT1 and INT2: carbanion/enolate intermediates (differs by a conformational rearrangement of the *β*-DMAM group). INT2 is also the reactant state (R2) for PT-2 (proton transfer from *β*-DMAM^+^ to C*α*). P: thioether adduct.

### Hydration of the various states along the reaction path

We analyzed the number of water within 3.5 Å of the Cys797 sulfur atom at different states along the reaction path using the aforementioned configurations extracted from the free energy simulations (Figure 4d). As expected, the charged Cys797 thiolate in the R1 state is well hydrated, with an average of 3.3 water molecules surrounding the sulfur atom; by contrast, the cysteine is desolvated in the neutral R0 state, with nearly no water. From the R1 to TS2 and INT1/INT2 states, the average hydration number decreases from 3.3 to 1.7 and 0.8/0.82, implying that the nucleophilic attack of Cys797 thiolate on the C*β* atom is coupled with the desolvation of the thiolate, which is consistent with a previous QM/MM study of the reaction between the covalent inhibitor 13-JAB with EGFR using the semiempirical DFTB/MM potential.^17^ The low hydration number remains from the INT1/INT2 states to the product state since the S-C covalent bond is formed.

## Concluding Discussion

The covalent bond formation mechanism between afatinib and Cys797 of the EGFR kinase was investigated using the combined PDDG-PM3/MM free energy simulations. Specifically, we evaluated two possible mechanisms, in which either the *β*-DMAM group on the acrylamide warhead or a nearby Asp800 acts as a general base to initiate the nucleophilic attack of the Cys797 thiolate. The calculated free energy profiles showed that in either mechanism, the reaction follows a classical Michael addition, which involves a stable carbanion/enolate intermediate and the nucleophilic attack as the rate-limiting step,^4,6–11,19^ consistent with the recent DFT QM/MM simulations of the ibrutinib reaction with the BTK kinase.^23^ However, the Michael addition mechanism of afatinib is in contrast to the concerted mechanism concluded from the earlier semiempirical DFTB3/MM simulations of the reactions of 13-JAB and osimertinib with EGFR.^17,18^ It is also inconsistent with the DFTB3/MM simulation derived mechanism of ibrutinib’s reaction with BTK, in which the carbonyl oxygen abstracts the proton from the cysteine and the enol-keto tautomerization is the rate-limiting step.^22^ We should note that both the inhibitor and protein in these studies^17,18,22^ are different from our work.

Our QM/MM simulations suggest that Mechanism 1 is more likely than Mechanism 2 (Asp800 is the general base/acid) for afatinib’s reaction with EGFR, as the former has a much lower barrier (19.5 vs. 31.3 kcal/mol) for the rate-limiting nucleophilic attack step, which is in close agreement with the barrier height of 21.6 kcal/mol estimated from the experimental rate constant of afatinib with EGFR.^26^ The interaction energy analysis for Mechanism 1 showed that Asp800 is the dominant contributor to the stabilization of the ion-pair (*β*-DMAM^+^/C797^−^) R1 state.

The simulations of Mechanism 1 show that the ion-pair state is a more stable reactant state than the neutral counterpart. Given that the *β*-DMAM group is charged in solution,^11^ this suggests that the deprotonation of Cys797, which is in the close proximity of the *β*-DMAM group in the crystal structure (sulfur-to-nitrogen distance is 3.6 Å, PDB 4G5J^16^), may be electrostatically driven, and the proton may be released to the solution water. Similarly, solution water may also protonate the carbanion (instead of PT-2 in Mechanism 1), resulting in a product state with a charged *β*-DMAM group. Note, we ruled out the involvement of a specific water molecule by examining the trajectory snapshots.

Regardless whether water is involved or not (i.e., variants of Mechanism 1), the NA step remains the same, and our data support the notion that the *β*-DMAM group enables the deprotonation of Cys797, while Asp800 plays a supporting role in stabilizing the reactant and transition states. These findings are in line with an early hypothesis regarding the role of *β*-methylamino substitution on acrylamides and the nearby Asp800.^27^ The role of Asp800 is also congruent with the solution simulation results, which showed that in the absence of Asp800 and the protein environ-ment, the PT-1 step is concerted with the NA step and the rate-limiting NA step has a much (10.3 kcal/mol) higher barrier.

The question of how protonated cysteines are activated in the protein environment to allow electrophilic additions is important for TCI design. Although the current study has delineated the role of the *β*-DMAM substitution, the question remains for the EGFR inhibitors without a similar substitution, e.g., osimertinib and 13-JAB. Is Asp800 the general acid/base in the absence of the *β*-DMAM substitution? Future work will address this question and the aforementioned hypothesis regarding the involvement of water. In this context, the recent GPU implementation of the all-atom particle mesh Ewald continuous constant pH MD technique^34^ may be used to determine the protonation state of the *β*-DMAM group in the enzyme environment. Notwithstanding, our work elucidates an important structure-activity relationship of acrylamides, aiding in the cysteine-directed TCI design.

## Computational Details

### Validation of the PDDG-PM3 QM potential

For the QM/MM simulations of afatinib reaction with EGFR, the QM region included the entire inhibitor and the side chain of Cys797 with the C*α* as the boundary atom (a total of 65 atoms, Figure 2b). For the solution model reaction simulations, the QM region included the entire inhibitor and methanethiol. To make such large QM regions computationally feasible, the semiempirical Pairwise Distance Directed Gaussian modified PM3 method (PDDG-PM3)^28,29^ was employed. The PDDG-PM3 calculated heats of formation of model compounds methanethiol, methylthiolate, and acrylamide (CH_2_**−**CHCONH_2_) are in good agreement with experimental data (Table S5).

We also compared the model reaction enthalpies calculated by the PDDG-PM3 method and the long-range corrected meta-GGA DFT method *ω*B97X-D3BJ,^35,36^ which was used in our previous work on the solution reactivities of model acrylamide warheads^11^ (Table S2).

The DFT calculations used the 6-311+G(d,p) basis set. PDDG-PM3 calculations were performed using CHARMM;^37^ *ω*B97X-D3BJ calculations were performed using the ORCA package (version 5.1.1).^38^

The QM/MM boundary atoms (C*α* of Cys797 in Mechanism 1 and 2, and C*α* of Asp800 in Mechanism 2) were treated by the generalized hybrid orbital (GHO) method.^39^ The GHO parameters for carbon atom in the combined PDDG-PM3/MM potential was obtained following the protocol used in Ref^40^ (See more details in SI).

### System preparation and MD simulations

For Mechanism 1, EGFR (except for the sidechain of Cys797), ions, and water comprised the MM region. The CHARMM22^41^ force field was used for the protein and ions, while the TIP3P model was used for water.^42^ The simulations of the reaction system was based on the X-ray co-crystal structure of EGFR in covalent complex with afatinib (PDB ID: 4G5J).^16^ The coordinates of the missing residues 721-724 and 747-756 were built based on the crystal structure of EGFR (PDB ID: 5U8L).^43^ The protonation states of titratable sites (Asp, Glu, His, Cys, and Lys) in EGFR were adopted from a previous study^13^ using the generalized-Born continuous constant pH MD titration simulations^44^ based on the crystal structure (PDB ID: 5U8L, ligand removed).^43^

The CHARMM package (version c45b2)^37^ was used for the simulations. The system (EGFR-afatinib complex and crystal water) was first minimized for 100 steps using the adopted-basis Newton Raphson (ABNR) method and then placed in a rectangular water box with at least 10 Å cushion between the protein heavy atoms and the edges of the box. Water molecules within 2.5 Å of any protein or inhibitor heavy atoms were deleted, and the crystal water molecules were kept. Three water in the bulk that were about 6 Å away from negatively charged residues on protein surface were selected and replaced with sodium ions to neutralize the system. In total, there were 26 crystal water and 12071 bulk water in the system. The solvated system was energy minimized using the PDDG-PM3/MM potential and the ABNR algorithm, first for the protein and QM atoms (40 steps), then water and ions (90 steps), finally the entire system (40 steps). Next, the system was heated to 298 K for 15 ps followed by equilibration for 50 ps with periodic boundary conditions and the constant NPT ensemble at 298 K and 1 atm. The Nośe-Hoover thermostat^45,46^ was used. The particle mesh Ewald method^47^ was employed for long-range electrostatic calculations with a real-space cutoff of 12 Å and a grid spacing of 1 Å. The non-bonded interactions were smoothly switched off between 12 and 13 Å. The nonbonded pair list was updated every 25 steps. All bonds involving hydrogen atoms were constrained by the SHAKE algorithm^48^ to allow an integration timestep of 2 fs.

For QM/MM simulations of the model reaction in solution, the structures of afatinib and methanethiol (built from the side chain of Cys797) were taken from the X-ray co-crystal structure (PDB ID: 4G5J).^16^ The system was solvated in a cubic water box (36x36x36 Å ^3^), and partitioned into the QM (afatinib and methanethiol) and MM (water) regions. The solvated system underwent energy minimization for 40 steps for the QM atoms, 60 steps for the MM water, and another 40 steps for the entire system. The heating and equilibration protocols were the same as for the protein system.

A similar system preparation and MD protocol was applied for Mechanism 2, except that the QM part includes 73 atoms (afatinib and the sidechains of Cys797 and Asp800) with two boundary atoms (C*α* of Cys797 and Asp800), and there were 12063 bulk water molecules and two sodium ions.

### Umbrella sampling free energy simulations

Following the MD equilibration, the free energy simulations were initiated from the EGFR-afatinib adduct state (or the methanethiol-afatinib adduct state for the model reaction) via the two possible mechanisms (Scheme 1). Free energy changes of the chemical steps were calculated in the reverse order of that shown in Scheme 1.

The adaptive umbrella sampling technique^49,50^ (module RXNCOR in CHARMM) was used. For the PT-2 step, 19 windows (23 for model reaction) in the reaction coordinate range of -1.8–1.8 Å (−1.6–2.4 Å for the model reaction). The NA and PT-1 steps were simulated through a two-dimensional free energy simulation with 327 windows (341 for the model reaction) in the reaction coordinate range of -2 to 3.2 Å for PT-1 and -2.2 to -0.3 Å for NA. For each umbrella sampling window, 100 ps equilibration was performed followed by 100 ps production run for trajectory collection. The convergence was verified by plotting the calculated free energies as a function of simulation time (Supplemental Figure S4). In the umbrella sampling, a biasing harmonic restraining potential with the force constant of 30–100 kcal/mol/Å ^2^ were placed on the reaction coordinates. 500 structures from each simulation window were saved for analysis. The weighted histogram analysis method (WHAM)^51^ was used to obtain the unbiased probability densities and free energy changes.

### QM/MM interaction energy decomposition analysis

A well-established energy decomposition method^32,33^ was applied to calculate the residue-based energy contributions to the stabilization or destabilization of the states along the reaction path. For the saved snapshots of a state, the QM/MM interaction energies of all residues in EGFR were calculated by sequentially zeroing the MM charges on a residue. The interaction energy between the residue and the QM region is defined as the difference in the QM electrostatic energy before and after the charge annihilation. The results were averaged over all the snapshots of each state.

## Acknowledgements

We thank Dr. Kwangho Nam for debugging the CHARMM QM/MM interface and the helpful discussion regarding QM/MM simulations. Financial support from the National Institutes of Health (R01CA256557) is acknowledged.

## Supporting Information Available

Protocol for the validation of the PDDG-PM3-GHO parameters. Protocol for the corrections of the PDDG-PM3 calculated energies based on the *ω*B97X-D3BJ DFT calculations. Protocol for selecting trajectory snapshots for the various states. Table S1 compares the calculated geometries of propane using the MM, PDDG-PM3/MM, PDDG-PM3, and HF/6-31G(d) methods. Table S2 contains the Mulliken atomic charges for atoms 1-5 of propane calculated by the PDDG-PM3 and PDDG-PM3/MM methods. Table S3 contains the modified parameters for the GHO carbon atom in combined PDDG-PM3/MM potential. Table S4 contains the calculated proton affinities for the blocked aspartic acid and cysteine. Table S5 compares the gas-phase heats of formation of model compounds from the PDDG-PM3 calculations and experiment. Table S6 and S7 contain the calculated reaction center distances and Mulliken atomic charges. Fig. S1 explains the QM/MM partition of propane. Fig. S2 and S3 compare the gas-phase model reaction enthalpies for Mechanism 1 and 2 from the PDDG-PM3 and DFT calculations. Fig. S4–S5 contain the calculated free energy profiles of the afatinib and model reactions according to Mechanism 1 without the DFT corrections. Fig. S6, S7 contain the calculated free energy profiles of the afatinib reaction in EGFR without (Fig. S6) and with (Fig. S7) the DFT corrections.

## Data Availability

The CHARMM MD program is freely available for academic use at https://academiccharmm.org/news/free-charmm.

The ORCA QM package is freely available for academic use at https://orcaforum.kofo.mpg.de. Cartesian coordinates of the model compounds are given in the downloadable PDF file (see Supporting Information). All simulation input files are available for download at https://github.com/JanaShenLab/Afatinib_reaction_mechanism.

## Abbreviations

DFT: density functional theory
DMAM: dimethylaminomethyl
EGFR: human epidermal growth factor receptor 1
MM: molecular mechanics
MD: molecular dynamics
TCI: targeted covalent inhibitor
QM: quantum mechanical

