## Supplemental information for "QM/MM simulations of EFGR with afatinib reveal the role of the *β*-dimethylaminomethyl substitution"

### Supporting Information: QM/MM simulations of EFGR with afatinib reveal the role of the $\beta$-dimethylaminomethyl substitution

Shuhua Ma,[*,†] Heeral Patel,[†] Craig A. Peeples,[‡] and Jana Shen[*,‡]

† *Department of Chemistry, Jess and Mildred Fisher College of Science and Mathematics, Towson University, Towson, MD 21252*

‡ *Department of Pharmaceutical Sciences, University of Maryland School of Pharmacy, Baltimore, MD 21201*

### Contents

**List of Tables** S-2

**List of Figures** S-2

### List of Tables

S1 Computed geometries of propane using MM, QM/MM, and QM methods . . S-8

S2 Atomic charges for the QM region calculated by the PDDG-PM3 and PDDG-PM3/MM methods . . . . . . . . . . . . . . . . . . . . . . . . . . . . . . S-9

S3 Modified parameters for the GHO carbon atom . . . . . . . . . . . . . . . . . S-9

S4 Calculated proton affinities of model cysteine and aspartic acid from the QM and QM/MM calculations . . . . . . . . . . . . . . . . . . . . . . . . . . S-9

S5 Model compound heats of formation from the PDDG-PM3 calculations and experiment . . . . . . . . . . . . . . . . . . . . . . . . . . . . . . . . . . S-9

S6 Reaction center distances in the different states of the reaction in EGFR and solution for Mechanism 1 . . . . . . . . . . . . . . . . . . . . . . . . . . S-10

S7 Reaction center atomic charges in the different states of the reaction in EGFR and solution for Mechanism 1 . . . . . . . . . . . . . . . . . . . . . . S-10

### List of Figures

S1 QM/MM partition scheme for propane . . . . . . . . . . . . . . . . . . . . . S-11

S2 Model reaction enthalpies in the gas phase calculated by the PDDG-PM3 and $\omega$B97X-D3BJ methods for Mechanism 1. . . . . . . . . . . . . . . . . . . S-11

S3 Model reaction enthalpies in the gas phase calculated by the PDDG-PM3 and $\omega$B97X-D3BJ methods for Mechanism 2. . . . . . . . . . . . . . . . . . . S-12

S4 **Convergence of the free energy calculations.** The calculated free energy of activation ($\Delta G^{\ddagger}$) and the reaction free energy ($\Delta G$) as a function of the simulation time (per umbrella window) for the PT-2 step of Mechanism 1 in EGFR. . . . . . . . . . . . . . . . . . . . . . . . . . . . . . . . . . . S-13

S5 Free energy profile of the afatinib reaction in EGFR for Mechanism 1 obtained from the free energy simulations **without the DFT correction**. . . . S-14

S6 Free energy profile of the model reaction in solution for Mechanism 1 obtained from the free energy simulations **without the DFT correction**. . . . S-14

S7 Free energy profile of the afatinib-EGFR reaction for Mechanism 2 obtained from the QM/MM free energy simulations **without the DFT correction**. . . S-15

S8 Free energy profile of the afatinib-EGFR reaction for Mechanism 2 **with the DFT corrections**. . . . . . . . . . . . . . . . . . . . . . . . . . . . . . . S-16

### Supplemental methods and protocols

#### Validation of the PDDG-PM3-GHO parameters for carbon

Following the previous protocol,[S1,S2] we tested the modified PDDG-PM3-GHO parameters for propane by comparing the geometries and Mulliken charge distributions calculated by the PDDG-PM3 method and PDDG-PM3/MM method. First, the propane molecule was optimized by the MM, PDDG-PM3, and HF/6-31G(d) methods. Then it was partitioned into QM and MM regions (Figure S1) and optimized by the PDDG-PM3/MM potential with C5 as the GHO boundary atom (with the $\beta_s$ value of -2.38200300 eV).

First, using the PDDG-PM3/MM optimized structure, the $\beta_p$ and $U_{pp}$ parameters were adjusted to reproduce the C4-C5 (C4 is a QM atom, C5 is the GHO boundary atom) bond length calculated from the full structure optimization by the PDDG-PM3 method (Table S1). Next, we made sure that the Mulliken atomic charge on C5 can reproduce the atomic partial charge in the CHARMM force filed (Table S2). The agreements were reached when $\beta_p$ and $U_{pp}$ were scaled by a factor of 1.26 and 0.969 (Table S3), respectively from the standard PDDG-PM3 parameters. All other bond lengths and bond angles calculated by the PDDG-PM3/MM method were either consistent with those calculated by the PDDG-PM3 method (for atoms 1-5) or with those calculated by MM method (for atoms 5-11). The largest bond length difference was 0.031 Å for the C5-C11 bond and the largest bond angle difference was 1.6° for the H6-C5-C11 angle between the PDDG-PM3/MM and MM calculated values. Similar to the performance of the PDDG-PM3 method compared to the HF/6-31G(d) optimized structure of propane, the PDDG-PM3/MM overestimates the C-H bonds (C4-H1, C4-H2, and C4-H3) by 0.008 Å and underestimates the C4-C5 bond lengths by 0.013 Å. The largest bond angle difference between the PDDG-PM3/MM and HF/6-31G(d) calculated values is 1.1° for C4-C5-H6(7). Overall, the adjusted $\beta_s$, $\beta_p$ and $U_{pp}$ values for the PDDG-PM3-GHO carbon atom allowed us to reproduce the geometries of propane.

In order to ensure the electronegativity of the GHO boundary carbon is in agreement with that of the carbon atom simulated by the standard PDDG-PM3 potential, we compared the Mulliken charges (Table S2) for atoms 1-5 of propane calculated by the PDDG-PM3-GHO/MM method with those calculated by the PDDG-PM3 method (for charges of atoms 1-4). The PDDG-PM3-GHO/MM calculated partial charges on atoms 1-4 agree well with those computed by the PDDG-PM3 method; and the PDDG-PM3-GHO/MM calculated partial charge on C5 is consistent with the assigned charge on this boundary atom in CHARMM force field.

Finally, the above PDDG-PM3-GHO parameters were used to test the proton affinities (Table S4) of cysteine (thiolate) and aspartic acid residues since their C$\alpha$ atoms were selected as the GHO boundary atoms in this work. We found that the proton affinities of cysteine (thiolate) and aspartic acid calculated by the PDDG-PM3/MM model deviates from those calculated from the standard PDDG-PM3 model by less that 1%.

#### Model reaction calculations and corrections based on the $\omega$B97X-D3BJ method

We calculated the reaction enthalpies for the model reactions of Mechanism 1 (Fig. S2) using the PDDG-PM3 method in comparison to the more accurate $\omega$B97X-D3BJ/6-311+G(d,p) method, which was recently used by us to study Michael additions with a large number of compounds.[S3] The PDDG-PM3 method was found to overestimate the reaction enthalpy of the PT-1 step by 10.8%, underestimates the reaction enthalpies of the NA and PT-2 steps by 2.7% and 14.9% respectively. Therefore, in the free energy profile (main text Scheme 2), the reaction free energy has been corrected by -0.7 kcal/mol (-6.3 kcal/mol to -7.0 kcal/mol, 10.8% decrease) for the PT-1 step (R0 to R1 state); and the 2.0 kcal/mol (-13.5 kcal/mol to -11.5 kcal/mol, 14.9% increase) for the PT-2 step (R2 to P state), the free energy profile of the reactions in EGFR for Mechanism 1 without corrections is shown in Figure S5. The reaction free energy for the NA step was not corrected due to a small difference between PDDG-PM3 and $\omega$B97X-D3BJ calculated reaction enthalpies. For reaction free energy in solution, only 5.2 kcal/mol (-34.7 kcal/mol in Figure S6 to -29.5 kcal/mol, a 14.9% increase) correction to the PT-2 step needed to be applied.

#### Validation of the PDDG-PM3 method

The reaction enthalpies for the model reactions of Mechanism 2 were similarly performed in the gas phase (Figure S3). The PDDG-PM3 method underestimates the reaction enthalpy of the PT-1 step by 145%, and overestimates the reaction enthalpies of the PT-2 step by 36.2%. The NA step model reaction is the same as in Mechanism 1. Considering these results, in the free energy profile (Figures S7 and S8) of Mechanism 2, the reaction free energy of the PT-1 step was corrected by 8.0 kcal/mol (0 kcal/mol to 8.0 kcal/mol, a 145% increase) and that of the PT-2 step was corrected by -10.8 kcal/mol (-29.7 kcal/mol to -40.5 kcal/mol, a 36.2% decrease).

#### Analysis of the states along the reaction path

The following reaction coordinate values were used to select trajectory snapshots for analyzing the different states along the reaction path for Mechanism 1. R0: (NA, PT-1) = (-2.15$\pm$0.2, -2.13 $\pm$0.2) Å; R1: (NA, PT-1) = (-2.05$\pm$0.2, 2.07$\pm$0.2) Å; TS2: (NA, PT-1) = (-0.88$\pm$0.12, 1.73$\pm$0.12) Å; INT1: (NA, PT-1) = (-0.30$\pm$0.2, 1.52$\pm$0.2) Å; INT2: (NA, PT-1) = (-0.30$\pm$0.2, 3.12 $\pm$0.2) Å; R2: PT-2 = -1.677$\pm$0.2 Å; TS4: PT-2 = -0.287 $\pm$0.1 Å; and P: PT-2 = 1.643$\pm$0.2 Å. Using these criteria, we saved 1178 snapshots for R0, 2301 for R1, 571 for TS2, 1712 for INT1, 1672 for INT2, 1517 for R2, 620 for TS4, and 1167 for P states.

For the simulations in solution, the states were similarly defined as follows. R0: (NA, PT-1) = (-2.21$\pm$0.2, -2.10$\pm$0.2) Å, 816 structures; TS2: (NA, PT-1) = (-0.98$\pm$0.12,

1.36$\pm$0.12) Å, 369 structures; INT: (NA, PT-1) = (-0.36$\pm$0.2, 1.59$\pm$0.2) Å, 1830 structures; R2: PT-2 = -1.543$\pm$0.2 Å, 1887 structures; TS4: PT-2 = -0.913$\pm$0.1 Å, 430 structures; and P: PT-2 = 2.617$\pm$0.2 Å, 440 structures.

### Supplemental tables

Table S1: Computed geometries of propane using MM, PDDG-PM3/MM, PDDG-PM3, and HF/6-31G(d) methods

| Bond length (Å) or angle (°) | MM | PDDG-PM3/MM | PDDG-PM3 | HF/6-31G(d) |
|---|---|---|---|---|
| $C_4-H_1$ | 1.111 | 1.095 | 1.096 | 1.087 |
| $C_4-H_2$ | 1.111 | 1.094 | 1.094 | 1.086 |
| $C_4-H_3$ | 1.110 | 1.094 | 1.097 | 1.087 |
| $C_4-C_5$ | 1.530 | 1.515 | 1.515 | 1.528 |
| $C_5-H_6$ | 1.115 | 1.125 | 1.109 | 1.087 |
| $C_5-H_7$ | 1.115 | 1.125 | 1.108 | 1.087 |
| $C_5-C_{11}$ | 1.530 | 1.561 | 1.515 | 1.528 |
| $C_{11}-H_8$ | 1.112 | 1.110 | 1.097 | 1.087 |
| $C_{11}-H_9$ | 1.112 | 1.109 | 1.095 | 1.086 |
| $C_{11}-H_{10}$ | 1.110 | 1.109 | 1.097 | 1.087 |
| $H_1-C_4-C_5$ | 110.4 | 111.6 | 111.7 | 111.1 |
| $H_2-C_4-C_5$ | 110.6 | 111.3 | 111.5 | 111.3 |
| $H_3-C_4-C_5$ | 110.5 | 111.6 | 111.7 | 111.1 |
| $C_4-C_5-C_{11}$ | 112.0 | 112.6 | 110.2 | 112.8 |
| $C_4-C_5-H_6$ | 109.3 | 110.5 | 110.2 | 109.4 |
| $C_4-C_5-H_7$ | 109.3 | 110.5 | 110.2 | 109.4 |
| $H_6-C_5-C_{11}$ | 109.3 | 107.7 | 110.2 | 109.4 |
| $H_7-C_5-C_{11}$ | 109.3 | 107.8 | 110.2 | 109.4 |
| $C_5-C_{11}-H_8$ | 110.5 | 110.0 | 111.7 | 111.1 |
| $C_5-C_{11}-H_9$ | 110.6 | 110.1 | 111.5 | 111.3 |
| $C_5-C_{11}-H_{10}$ | 110.5 | 110.0 | 111.7 | 111.1 |

In the PDDG-PM3/MM calculations, atoms 1-5 were treated by the PDDG-PM3 potential, atoms 6-11 were treated molecular mechanically, and C5 was the GHO boundary atom.

Table S2: Mulliken atomic charges for atoms 1-5 of propane calculated by the PDDG-PM3 and PDDG-PM3/MM methods).

| Atoms | PDDG-PM3 | PDDG-PM3/MM |
|---|---|---|
| $H_1$ | 0.09 | 0.09 |
| $H_2$ | 0.09 | 0.09 |
| $H_3$ | 0.09 | 0.09 |
| $C_4$ | -0.27 | -0.27 |
| $C_5$ | -0.18* | -0.18 |

Charge unit is atomic unit. *The charge on the QM/MM boundary atom C5 is the MM charge.

Table S3: Modified parameters (eV) for the GHO carbon atom in combined PDDG-PM3/MM potential.

| | PDDG-PM3 | PDDG-PM3-GHO |
|---|---|---|
| $\beta_s$ | -11.952818190434 | -2.38200300 |
| $\beta_p$ | -9.9224112120852 | -12.50223813 |
| $U_{pp}$ | -36.461255999939 | -35.33095706 |

All other QM parameters for the GHO carbon were taken from the original PDDG-PM3 parameter set.

Table S4: Calculated proton affinities for blocked model cysteine and aspartic residues in the gas phase using PDDG-PM3 and PDDG-PM3/MM methods

| | PDDG-PM3 | PDDG-PM3/MM |
|---|---|---|
| Cys | 327.62 | 328.08 |
| Asp | 346.05 | 342.67 |

The unit is kcal/mol. The experimental value of 365.7 kcal/mol was used as the heat of formation of proton.

Table S5: Comparison of the heats of formation of model compounds in the gas phase (kcal/mol) from the PDDG-PM3 calculations and experiment

| | PDDG-PM3 | Experiment | Reference |
|---|---|---|---|
| $CH_3SH$ | -5.3 | -5.46 | Ref[S4] |
| $CH_3S^-$ | -19.2 | -14.3 | Ref[S5] |
| $CH_2{=}CHCONH_2$ | -29.8 | -31.12 | Ref[S6] |

Table S6: Selected distances (Å) at the reaction center in the different states along the reaction path in EGFR and solution for Mechanism 1

| Distance | R0 | R1 | TS2 | INT1 | INT2/R2 | TS4 | P |
|---|---|---|---|---|---|---|---|
| S–C$\beta$ | 3.42 | 3.36 | 2.28 | 1.85 | 1.85/1.88 | 1.84 | 1.85 |
| Solution | 3.45 | n/a | 2.34 | 1.85 | 1.85 | 1.85 | 1.84 |
| C$\alpha$–C$\beta$ | 1.34 | 1.34 | 1.39 | 1.48 | 1.47/1.46 | 1/49 | 1.53 |
| Solution | 1.34 | n/a | 1.38 | 1.47 | 1.47 | 1.48 | 1.52 |
| C–C$\alpha$ | 1.48 | 1.49 | 1.44 | 1.39 | 1.39/1.39 | 1.45 | 1.52 |
| Solution | 1.48 | n/a | 1.46 | 1.39 | 1.39 | 1.41 | 1.53 |
| C–O | 1.23 | 1.25 | 1.25 | 1.27 | 1.27/1.27 | 1.25 | 1.24 |
| Solution | 1.24 | n/a | 1.25 | 1.27 | 1.27 | 1.27 | 1.24 |

The distances are in Å. The values from the model reactions in solution are included in the rows starting with Solution.

Table S7: Mulliken charges of the selected atoms of the reaction center in the different states along the reaction path in EGFR and solution for Mechanism 1

| Atom | R0 | R1 | TS2 | INT1 | INT2 | R2 | TS4 | P |
|---|---|---|---|---|---|---|---|---|
| C$\alpha$ | -0.30 | -0.24 | -0.60 | -0.70 | -0.70 | -0.73 | -0.64 | -0.22 |
| solution | -0.27 | | -0.49 | -0.73 | | -0.76 | -0.75 | -0.21 |
| C$\beta$ | -0.06 | -0.14 | 0.02 | -0.12 | -0.11 | -0.09 | -0.19 | -0.26 |
| solution | -0.12 | | -0.02 | -0.09 | | -0.10 | -0.12 | -0.25 |
| C | 0.29 | 0.25 | 0.30 | 0.29 | 0.27 | 0.28 | 0.29 | 0.24 |
| solution | 0.26 | | 0.31 | 0.31 | | 0.31 | 0.31 | 0.22 |
| O | -0.39 | -0.33 | -0.46 | -0.55 | -0.55 | -0.52 | -0.43 | -0.40 |
| solution | -0.35 | | -0.41 | -0.50 | | -0.50 | -0.48 | -0.37 |

The charges are in atomic unit. The values from the model reactions in solution are included in the rows starting with Solution.

### Supplemental figures

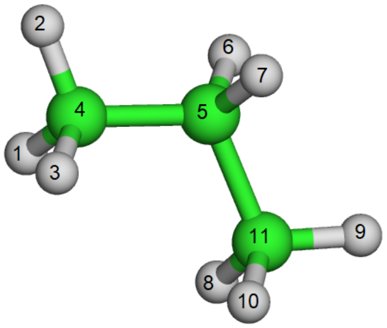

Figure S1: Structure of propane. In the combined PDDG-PM3/MM calculations, atoms 1-5 were treated quantum mechanically by the PDDG-PM3 potential, C5 was the GHO boundary atom, and atoms 6-11 were treated molecular mechanically in CHARMM.

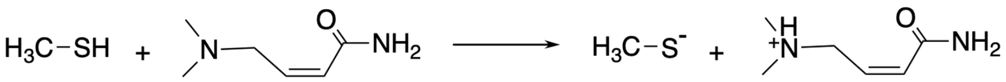

**1 (PT-1)**

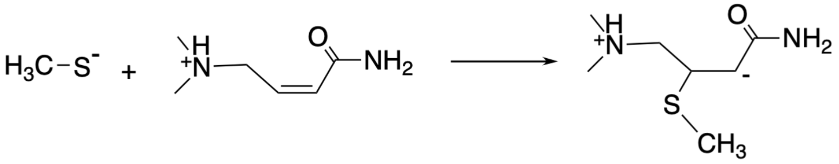

**2 (NA)**

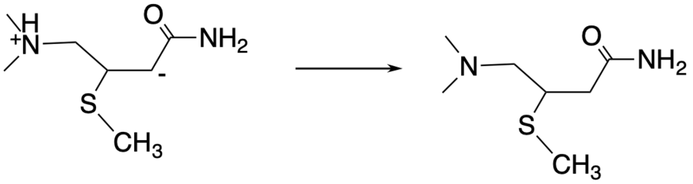

**3 (PT-2)**

| Reaction | PDDG/PM3 | ωB97X-D3BJ | %diff |
|---|---|---|---|
| **1 (PT-1)** | 145.1 | 129.4 | +10.8% |
| **2 (NA)** | -104.0 | -101.2 | -2.7% |
| **3 (PT-2)** | -60.3 | -51.3 | -14.9% |

Figure S2: Model reaction enthalpies in the gas phase calculated by the PDDG-PM3 and ωB97X-D3BJ methods for Mechanism 1.

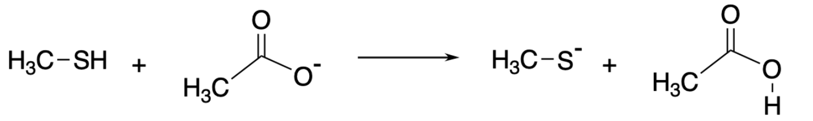

**1 (PT-1)**

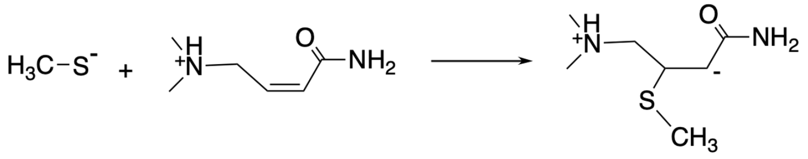

**2 (NA)**

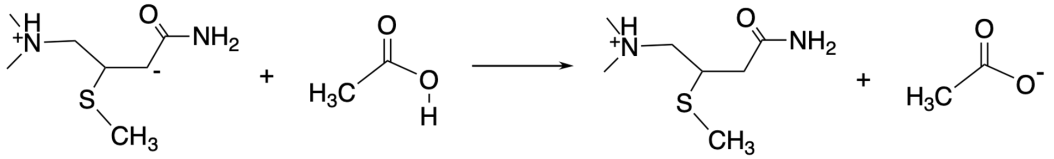

**3 (PT-2)**

| Reaction | PDDG/PM3 | ωB97X-D3BJ | Expt.* | %diff |
|---|---|---|---|---|
| **1 (PT-1)** | 5.5 | 15.9 | 13.5 | -145% |
| **2 (NA)** | -104.0 | -101.2 | | -2.7% |
| **3 (PT-2)** | 72.1 | 46.0 | | +36.2% |

*cis- acetic acid conformation

Figure S3: Model reaction enthalpies in the gas phase calculated by the PDDG-PM3 and $\omega$B97X-D3BJ methods for Mechanism 2.

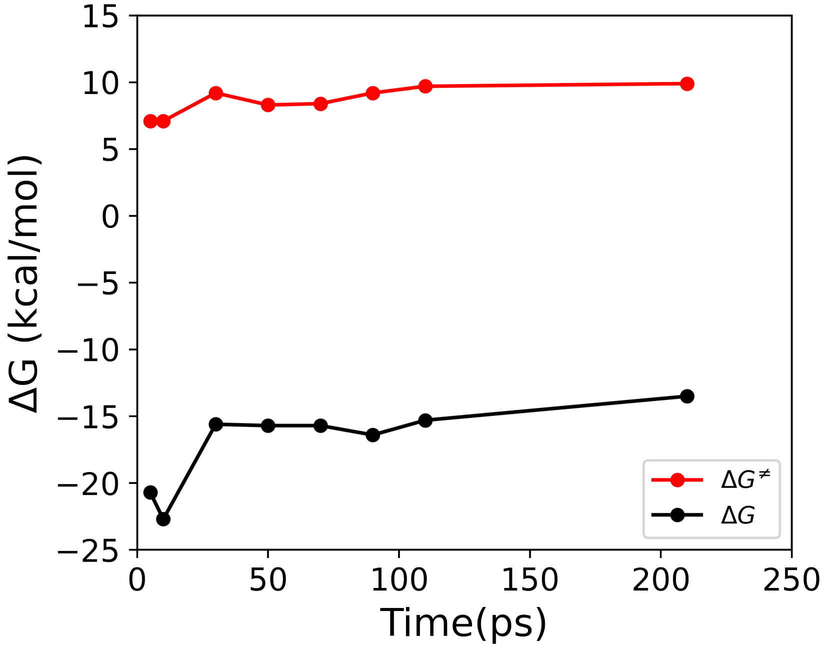


Figure S4: **Convergence of the free energy calculations.** The calculated free energy of activation ($\Delta G^{\ddagger}$) and the reaction free energy ($\Delta G$) as a function of the simulation time (per umbrella window) for the PT-2 step of Mechanism 1 in EGFR.

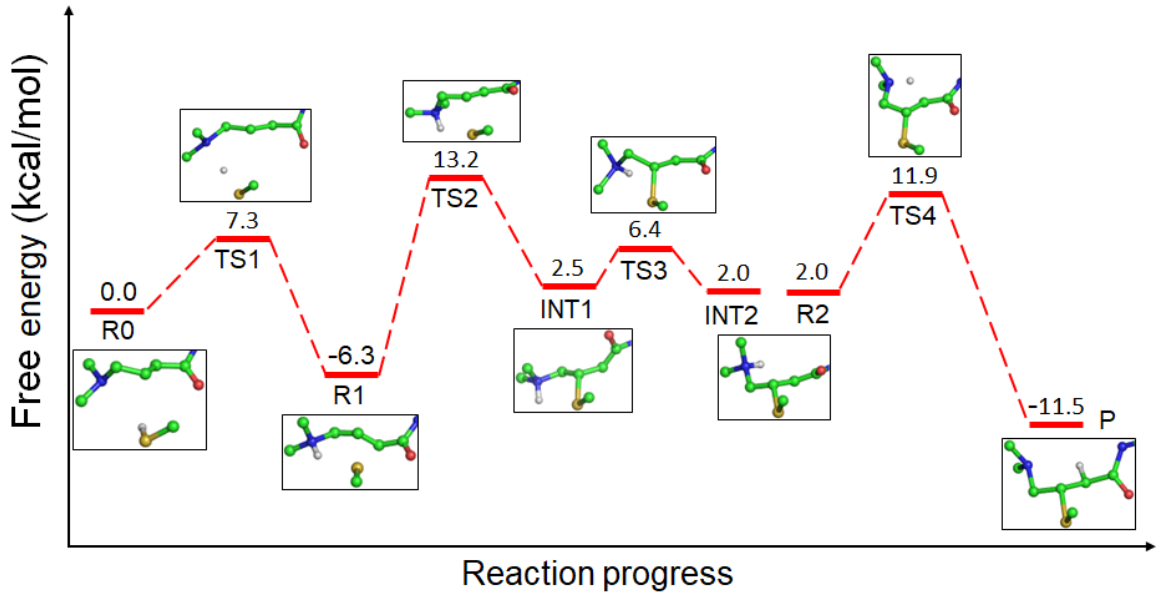


Figure S5: Free energy profile of the afatinib reaction in EGFR for Mechanism 1 obtained from the free energy simulations **without the DFT correction**.

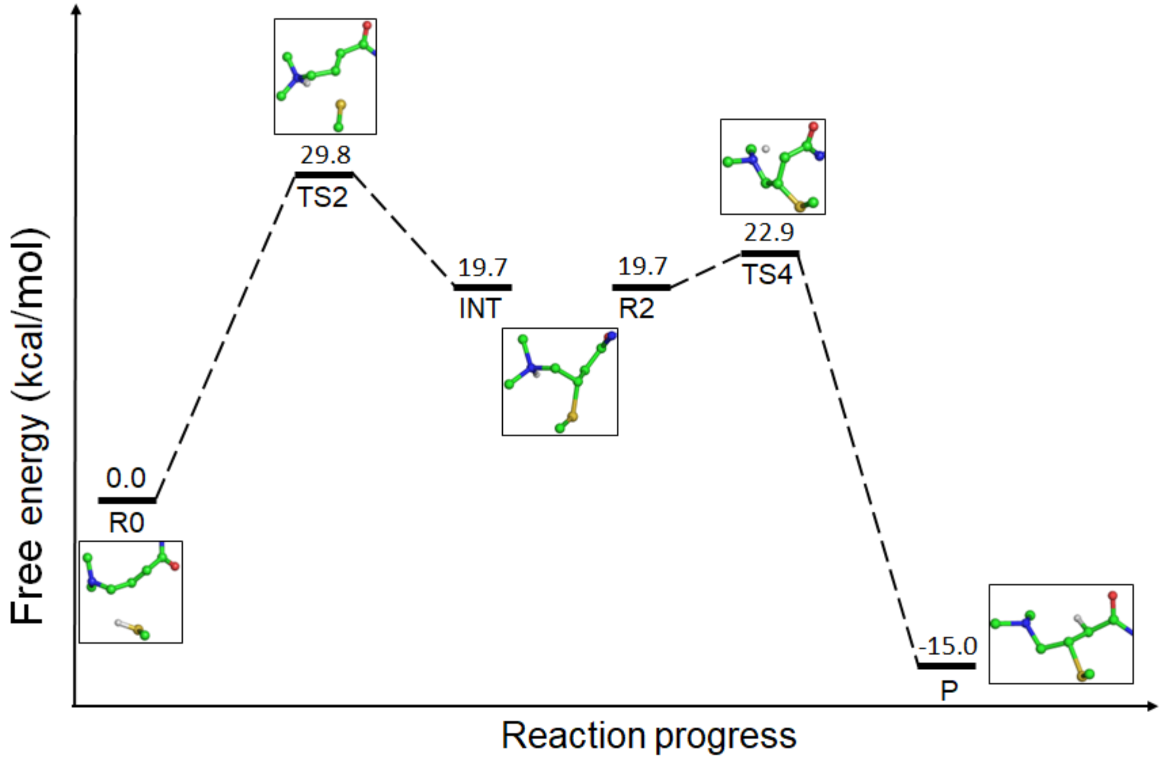


Figure S6: Free energy profile of the model reaction in solution for Mechanism 1 obtained from the free energy simulations **without the DFT correction**.

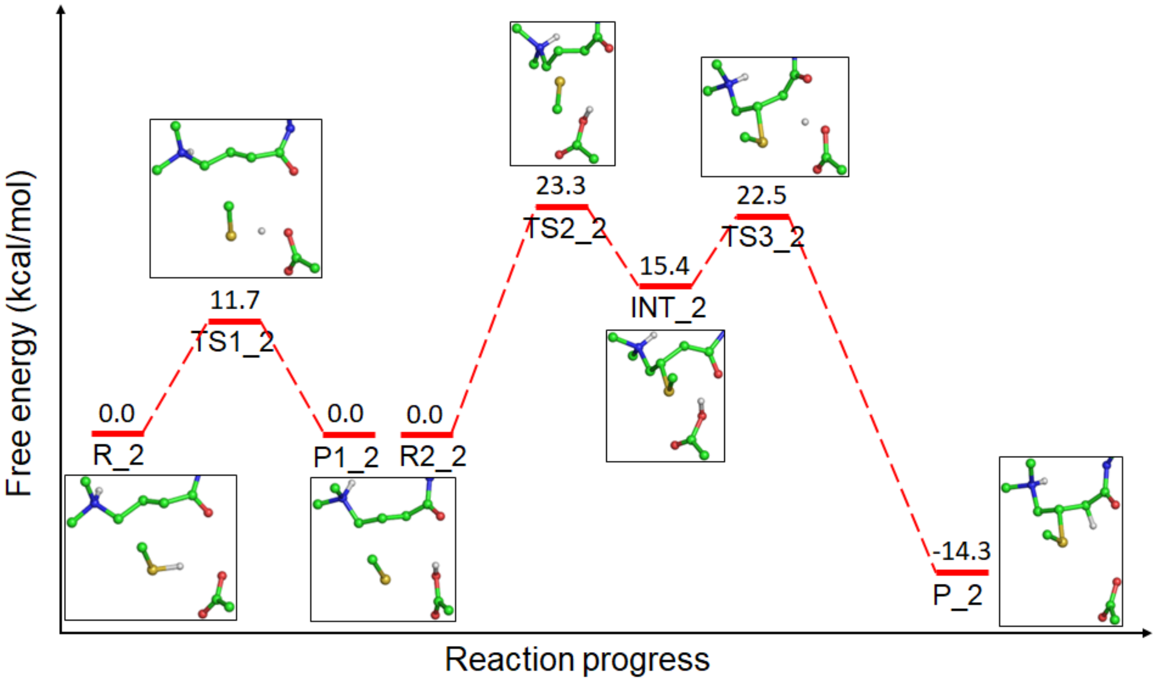


Figure S7: Free energy profile of the afatinib-EGFR reaction for Mechanism 2 obtained from the QM/MM free energy simulations **without the DFT correction**.

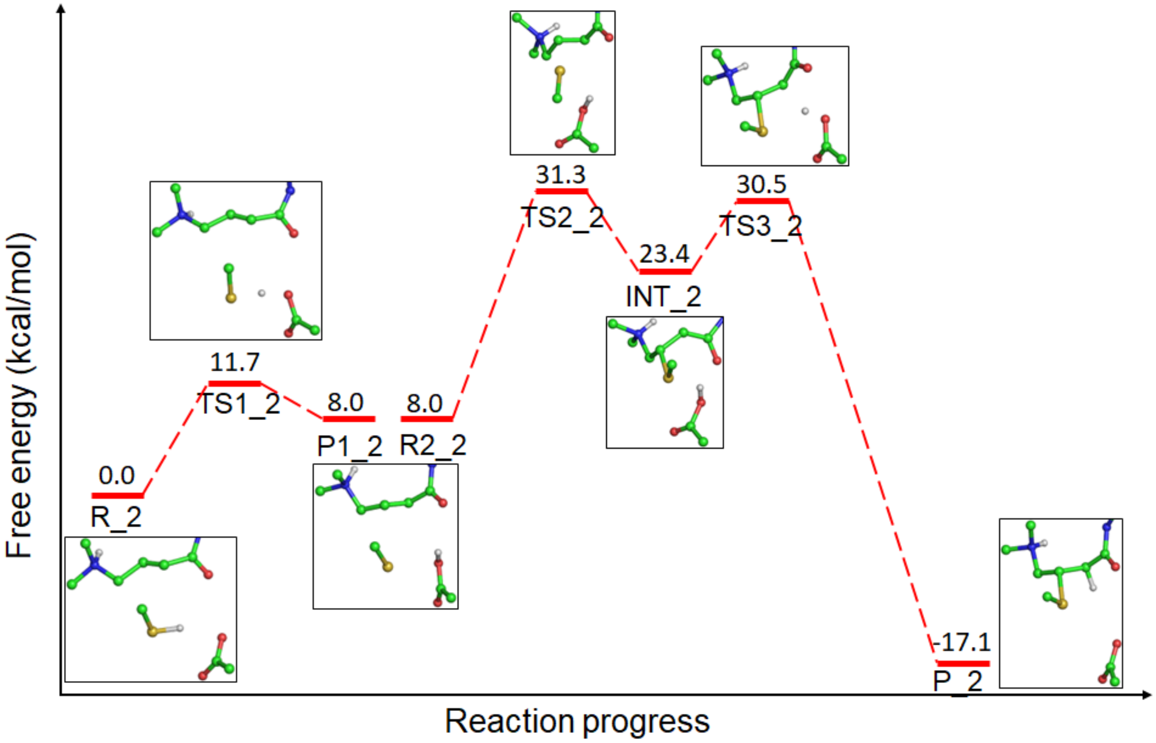


Figure S8: Free energy profile of the afatinib-EGFR reaction for Mechanism 2 **with the DFT corrections**.

(S6) Steele, W. V.; Chirico, R. D.; Nguyen, A.; Hossenlopp, I. A.; Smith, N. K. Determination of some pure compound ideal-gas enthalpies of formation.
